# ConstrastivePose: A contrastive learning approach for self-supervised feature engineering for pose estimation and behavorial classification of interacting animals

**DOI:** 10.1101/2022.11.09.515746

**Authors:** Tianxun Zhou, Calvin Chee Hoe Cheah, Eunice Wei Mun Chin, Jie Chen, Hui Jia Farm, Eyleen Lay Keow Goh, Keng Hwee Chiam

## Abstract

In recent years, supervised machine learning models trained on videos of animals with pose estimation data and behavior labels have been used for automated behavior classification. Applications include, for example, automated detection of neurological diseases in animal models. However, there are two problems with these supervised learning models. First, such models require a large amount of labeled data but the labeling of behaviors frame by frame is a laborious manual process that is not easily scalable. Second, such methods rely on handcrafted features obtained from pose estimation data that are usually designed empirically. In this paper, we propose to overcome these two problems using contrastive learning for self-supervised feature engineering on pose estimation data. Our approach allows the use of unlabeled videos to learn feature representations and reduce the need for handcrafting of higher-level features from pose positions. We show that this approach to feature representation can achieve better classification performance compared to handcrafted features alone, and that the performance improvement is due to contrastive learning on unlabeled data rather than the neural network architecture.

**Author Summary:** Animal models are widely used in medicine to study diseases. For example, the study of social interactions between animals such as mice are used to investigate changes in social behaviors in neurological diseases. The process of manually annotating animal behaviors from videos is slow and tedious. To solve this problem, machine learning approaches to automate the video annotation process have become more popular. Many of the recent machine learning approaches are built on the advances in pose-estimation technology which enables accurate localization of key points of the animals. However, manual labeling of behaviors frame by frame for the training set is still a bottleneck that is not scalable. Also, existing methods rely on handcrafted feature engineering from pose estimation data. In this study, we propose ConstrastivePose, an approach using contrastive learning to learn feature representation from unlabeled data. We demonstrate the improved performance using the features learnt by our method versus handcrafted features for supervised learning. This approach can be helpful for work seeking to build supervised behavior classification models where behavior labelled videos are scarce.

## Introduction

Analysis of animal behavior is critical in the field of neuroscience to study brain function, and crucial for the assessment of treatment efficacy in preclinical testing. With the advancement of molecular tools for intervention in animal models, accurate and efficient detection and quantification of animal behavior is increasingly sought after. While human annotators remain the gold standard in behavior scoring, they can get fatigued or overwhelmed by the vast number of behaviors to score, in addition to the complexity of differentiating specific behaviors. It takes about 22 man-hours to annotate a one-hour video by frame with high confidence[1]. Other problems with human annotation are the difficulty of ensuring high quality of annotation due to well documented factors such as variability between different annotators, observer bias and observer drift[1–4].

Automated video analysis has been introduced to help allow a semi-high throughput workflow for behavioral screening in research[5]. Commercial behavior tracking software packages (e.g. EthoVision, ANY-maze), or those that are incorporated in the behavioral assay equipment hardware (e.g. Med Associates Inc., Campden Instruments Ltd.,) are often costly, and have low customizability to user-specific experimental setting. Additionally, some studies have shown that many commercial software lack sensitivity due to poor animal tracking and are unable to dissociate complex animal behaviors[5–9]. Due to such drawback, machine learning-based approaches using open-source software and videos acquired with consumer grade cameras have steadily being embraced by animal behavioral scientists for automated tracking and analysis of complex behaviors in their research models. For example, Wu et al.[10] developed a machine-learning image-analysis program that automatically tracks leg claw positions of freely moving flies recorded on high-speed video, producing a series of gait measurements. Their fully automated leg tracking of *Drosophila* neurodegeneration models reveals distinct conserved movement signatures. Hong et al.[11] studied interactions of mice with gene mutations associated with autism using machine learning based video tracking and classification and detected social interaction deficits compared to those without the mutations. Van den Boom et al.[6] applied open-source machine learning classification software to study SAPAP3 knockout mice and confirmed that they engage in more grooming than wildtype mice from the same litter both in number of bouts and grooming duration.

The common workflow[1,5] for machine learning animal behavior classification is to first extract features from the video, commonly in the form of pose estimation. Feature engineering is then performed by computing hand-crafted features, such as animal orientation and length, from animal pose estimation. Finally, a machine learning algorithm, either supervised learning or unsupervised learning, is applied on those features. In supervised learning, the classifier is trained on the features and behavior labels, and the trained classifier can be used to classify behaviors in new videos. We focus on the supervised learning workflow as it is more commonly used in literature.

There are two weaknesses of this typical workflow for practitioners. Firstly, the requirement of creating a large labeled training set for the machine learning model to achieve good classification accuracy. E.g., in the supervised classification of mice behavior, up to 260 minutes, and 135 minutes of video were annotated in [12], and [13], requiring approximately 95 and 50 man-hours of work respectively to build the training and validation sets. Secondly, engineering handcrafted features from pose-estimation data relies on experience and trial-and-errors. A summary of various feature engineering approaches found in literature is provided in S1 Table.

In this study, we develop a method, which we refer to as ConstrastivePose, that seeks to address these two weaknesses. The ConstrastivePose method is trained using contrastive learning on unlabeled data, and then fine-tuned with a small amount of labeled data.

Contrastive learning, a form of self-supervised learning that learns useful representations of data for classification without the need for labels. It is trained by *contrasting* similar data against dissimilar data. For a given datapoint in a training batch, similar data is generated by data augmentation of itself while dissimilar data are simply other datapoints in the training batch. Through contrastive learning, ConstrastivePose leverages the availability of large sets of unlabeled data generated with the automated and easily scalable pose estimation data generation process. The data representation learnt by ConstrastivePose also reduces the need for manual feature engineering from pose estimation. With fine-tuning after contrastive learning, ConstrastivePose achieves better performance on downstream supervised learning task than handcrafted feature engineering. Through this work, we hope to improve behavior classification performance and alleviate the reliance on manual annotations by trained behavioral scientists to decipher animal behavior.

## Results

### ConstrastivePose learns features that exhibit similar structure as handcrafted features

ConstrastivePose uses contrastive learning to reduce differences in representation between a set of pose estimation and its random augmented version and enlarges their differences with other examples in the batch. We trained a neural network to take in the pose-estimation and output an embedding which can be interpreted as features constructed by the neural network from the original data. To demonstrate how this works, we first apply ConstrastivePose to the Caltech Mouse Social Interactions (CalMS21) Dataset[14] Task 1, which contains videos of two mice interacting that have been labeled for key body part positions and one of four behaviors: investigation, attack, mount and others (more details provided in materials and methods section). Visualization of the embedding space with UMAP for the CalMS21 dataset in **Fig *1*** showed that contrastive learning was able to learn a representation that is similar to the embedding spaces formed by handcrafted feature engineering methods, as opposed to the original feature representation with no feature engineering. We can see that in **Fig *1*** panel *a*, the original feature representation does not show any coherent groups or clusters between different behaviors. In panel *b*, the representation of handcrafted features shows distinction between interacting behaviors (investigation, attack and mount) and non-interacting behaviors (others). The learnt representation in panel *c* was able to achieve similar results as in panel B, with clearer distinction and more separable structure.

**Fig 1.**
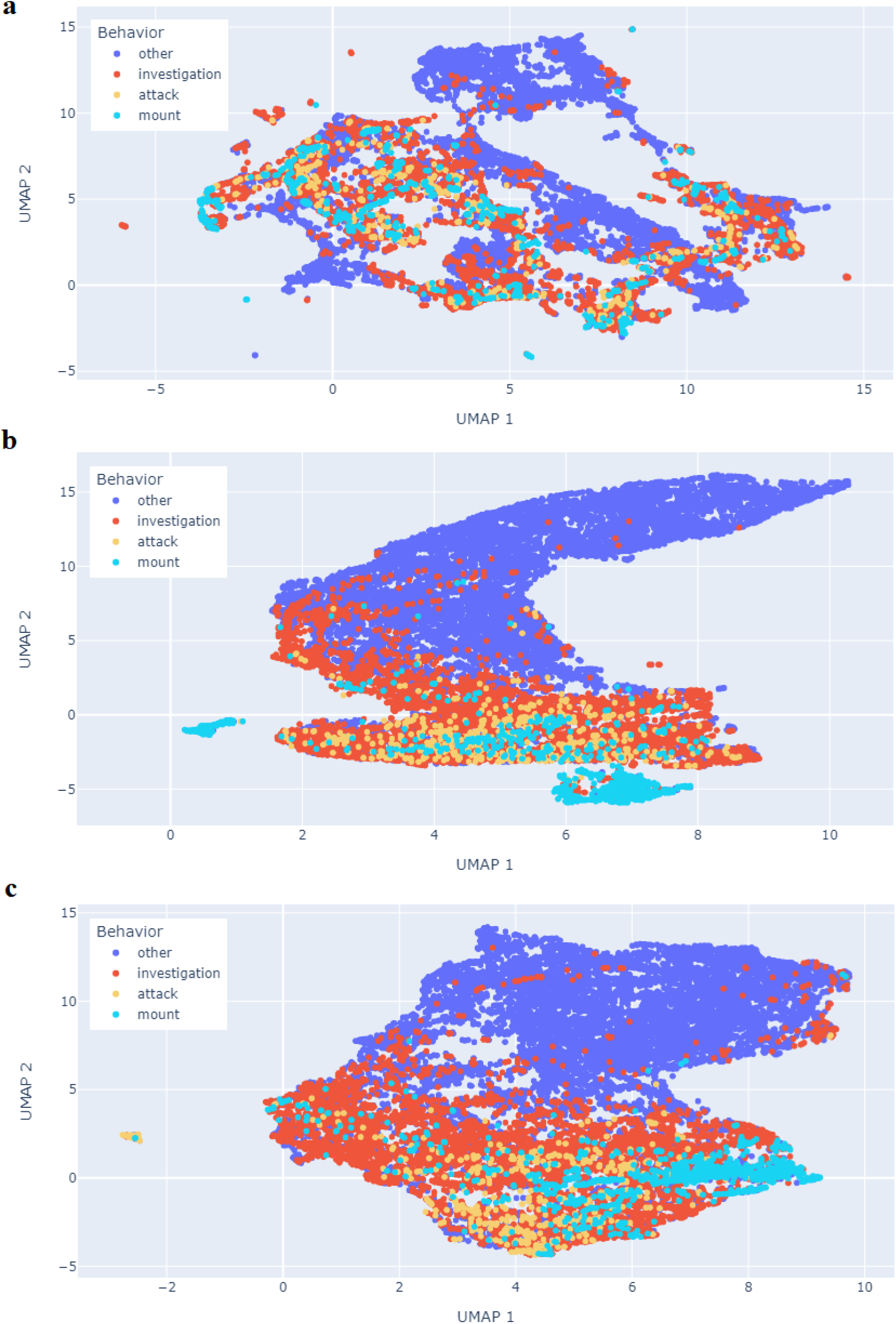
Visualization of feature representations for CalMS21 dataset using UMAP. **(a)** Representation of the original untransformed features. **(b)** Representation of the handcrafted features with the best classification performance in Table 3. **(c)** Representation of learnt features through contrastive learning. The learnt representation in panel c was able to achieve similar results as in panel b, with clearer distinction and more separable structure.

### ConstrastivePose outperforms no feature engineering, and is on-par with handcrafted feature engineering for supervised learning

To test how well the feature representations learned by ConstrastivePose performs on supervised learning, we compared our method against handcrafted engineered features that were commonly used in literature (S1 Table). For our method, we trained ConstrastivePose on unlabeled data and then fine-tuned it on a small set of labeled data.

A random forest model was employed to compare the test performances of the different feature engineering methods. Tree based ensemble methods such as random forest are easy to train and are one of the most popular supervised learning methods used in animal behavior supervised classification[1]. Each of the random forest model is trained on a separate set of engineered features as inputs and then tested with unseen test data.

The performance of the models trained on different combinations of engineered features are summarized in **Table 1**. Macro-averaging was used for the metrics because of class imbalance to treat all classes as equally important and avoid overoptimistic estimation of the classifier performance due to the majority class. We found similar or higher scores for precision, recall, F1, and accuracy for our method compared to supervised learning. In particular, the macro F1 score for our method was at least 0.05 higher than the next highest supervised engineering feature, indicating greater multiclass classification performance. (Complete classification results for each class are provided in S7)

**Table 1.**
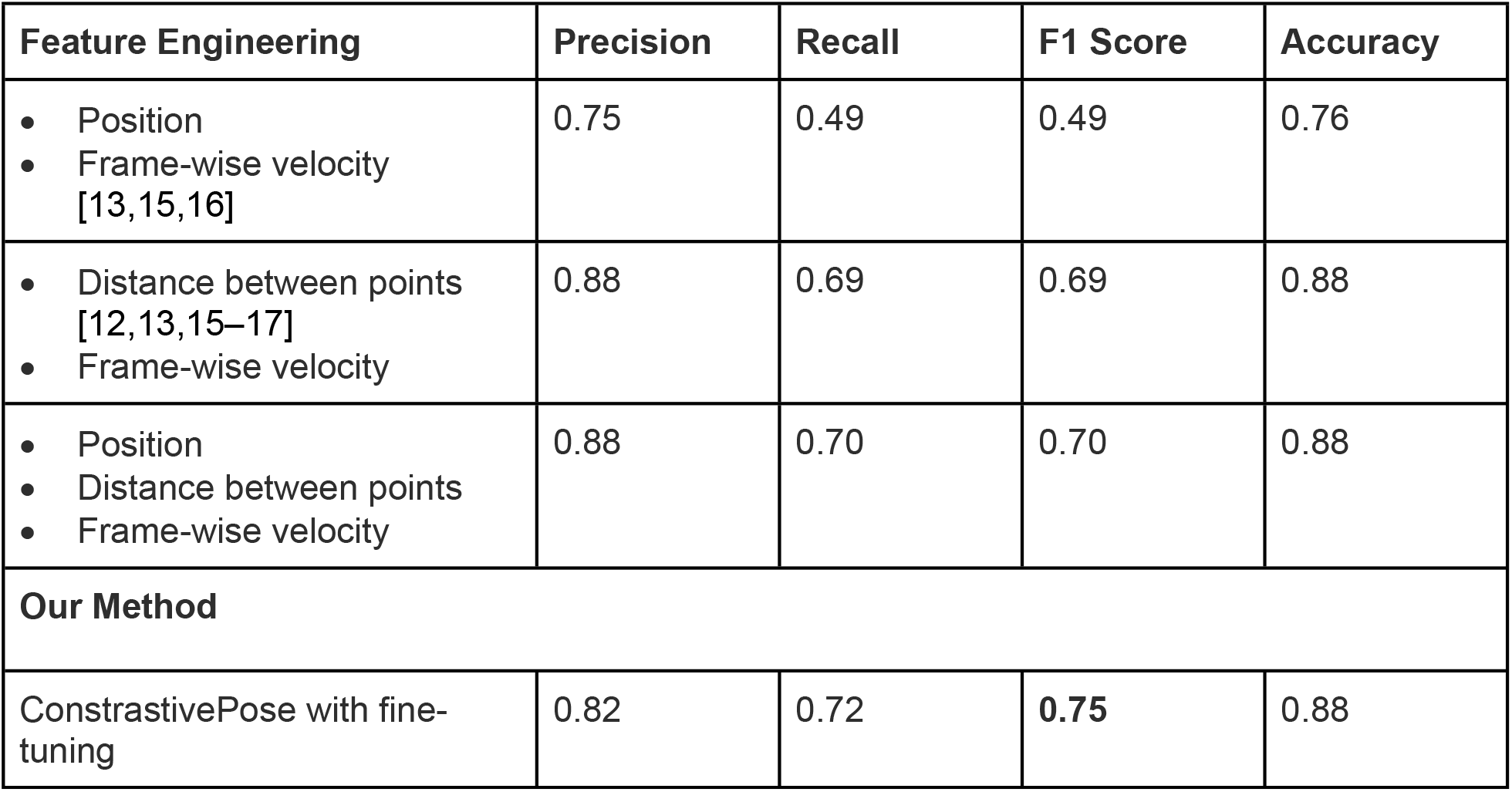
Comparison of classification performance for various handcrafted feature engineering methods for CalMS21 dataset.

To further validate the performance of our model, we applied it to a new set of data that we generated from our in-house experiments. Two wild type mice were housed in a cage and videos of them interacting were captured. Either animal can be behaving individually, such as self-grooming, or one can be following the other, or engaging in sniffing the body or the anogenital region of the other, or both animals can come together and perform nose-to-nose sniffing. (See S2 for list of behaviors.) Thus, this dataset is more challenging as it contained more than twice as many behavior classes compared to CalMS21. Furthermore, the mice in the videos were of the same color and size, which made it difficult for pose-estimation software to extract pose with high accuracy. Hence, the pose estimation input was noisy and contained some missing or erroneous data. This is the case for both DeepLabCut[4], a popular pose estimation software, as well as using YOLO-based object detection algorithm[18], suggesting the inherent difficulty of the video rather than an issue with the pose estimation software choice. Nevertheless, despite these challenges, ContrastivePose provides similar performance advantages on this dataset. The results are summarized in **Table 2**.

**Table 2.** Comparison of classification performance for various handcrafted feature engineering methods for in-house experiments.

| Feature Engineering | Precision | Recall | F1 Score | Accuracy |
| --- | --- | --- | --- | --- |
| Position of corners<br>Distance between points<br>Frame-wise velocity | 0.15 | 0.17 | 0.11 | 0.35 |
| Position of corners<br>Distance between points<br>Frame-wise velocity<br>Overlap in bounding boxes | 0.24 | 0.39 | 0.25 | 0.65 |
| **Our Method** |  |  |  |  |
| ContrastivePose with fine-tuning | 0.30 | 0.44 | **0.32** | 0.72 |

Well-designed handcrafted features, in this case the overlap between bounding boxes, can have good prediction power. The overlap in bounding boxes is computed using the intersection area between two bounding boxes. When interacting animals come into close contact with each other, bounding boxes of the body parts will tend to overlap and the intersection area provide information about the type and extent of contact between a pair of body parts. As seen in **Table 2**, when overlap in bounding box feature was added, the classification scores increased substantially. By supplementing the features learnt by ContrastivePose with well-designed handcrafted features like the overlap between bounding boxes, which is easily done, it can achieve better performance than just the handcrafted feature set measurably.

## Discussions

Machine learning methods for animal behavior classification typically follow a two-step process: feature extraction from the video, commonly in the form of pose estimation, followed by machine learning classification. In recent years, pose-estimation or pose-tracking has advanced rapidly with the introduction of deep learning methods in computer vision that allows for markerless tracking of various user-selected body parts of animals to be accurately tracked in video. Open-source tools such as DeepLabCut[4], DeepPoseKit[19] and YOLO[18,20] are now popular and widely used among researchers. We focus on pose estimation features as input due to their popularity. In most cases, hand-crafted features such as animal orientation and length are then computed from the pose-estimation to be used in the second step.

In the second step, machine learning is used to classify behaviors using the features extracted and computed in the first step. Machine learning methods generally fall under supervised and unsupervised learning[1]. Supervised learning trains a model with true labels provided. There have been many works using supervised learning methods such as random forest[12,15], support vector machines (SVMs)[21], and neural networks[22]. Unsupervised learning seeks to discover inherent structure within the data, typically by finding various spatial groupings in a feature space after some form of dimensionality reduction. These spatial groupings may correspond to various human defined behaviors or behavior “motifs” upon inspection[1]. However, user oversight is still necessary at the end to ensure accuracy and explainability of output variables.

A main weakness of supervised classification of behaviors from pose-estimation is the requirement of accurate annotation for the creation of labels needed for training, which currently relies on human input. Supervised learning is known to perform better with more available labeled data. However, as mentioned previously, creating high quality manual labeling is a time-consuming process. Generating more training data will require more man-hours spent on labeling.

Moreover, the use of multiple human labelers or even the same labeler on different working sessions inevitably introduces variability due to observer bias and observer drift.

Another aspect of most existing machine learning workflows for animal behavior classification using pose-estimation data as input, is feature engineering. Various methods in literature mostly rely on handcrafted features computed from the pose-estimation data (S1 Table). Feature engineering from pose-estimation data can be a tedious and difficult process. Handcrafting features depend much on the intuition and experience of the designer. In this process, poorly designed feature engineering can potentially fail to capture necessary information and relationships that are needed to obtain high classification accuracy.

To overcome the burden of tedious manual annotation of behavior from videos and reliance on trained observers, we developed ConstrastivePose, which uses contrastive learning to train on pose estimation data alone, and output behavior classifications. This method reduces the need for feature engineering and is able to learn from large unlabeled datasets to improve the model learnt representation. Contrastive learning was first successfully applied in computer vision to leverage the fact that there are huge amounts of unlabeled images available compared to labeled images. Through self-supervised learning on a larger set of unlabeled images, and then fine-tuning the representation learnt for downstream tasks like image classification and object detection, it is possible to obtain quality performance with much lesser labeled data (18–20). Our method, ConstrastivePose, is a novel application of contrastive learning on the problem of classifying animal behavior from pose estimation data.

ConstrastivePose has two main advantages over existing methods. Firstly, this approach enables the leveraging of larger amounts of pose-estimation extracted from unlabeled video to improve predictions, alleviating the bottleneck of lesser available labeled video data. Secondly, this approach reduces the need for feature engineering from user-defined to learning from the data itself. In comparison, current methods with engineered features are static in the sense that once the user defines the rules or calculations to generate the features, e.g. pairwise distance, angle between subjects, these rules are fixed no matter how much pose-estimation data is available. The self-supervised learning approach, however, can leverage on more available data to improve its feature extraction ability. In the results section, we have shown that by performing contrastive learning on a larger set of unlabeled pose-estimation data, and then fine-tuning with a small set of labeled training data, ConstrastivePose can achieve better performance on downstream supervised classification than using handcrafted features.

We also trained a model with the same neural network architecture using a small set of labeled training data alone, without the contrastive pre-training, as a feature extractor to understand if the performance improvement was due to the use of contrastive learning or simply the strength of the neural network architecture itself as a feature extractor (Refer to experiment set-up details in S3 Fig *1*). The results summarized in **Table 3** show that training from scratch on labeled data alone performs worse than training with contrastive learning. This demonstrates that the performance improvement comes from representation learnt during contrastive learning, and that the method is an effective way of boosting performance by incorporating information from unlabeled pose-estimation data.

**Table 3.** Comparison of classification performance for ConstrastivePose and neural network without pretraining.

| Method | Precision | Recall | F1 Score | Accuracy |
| --- | --- | --- | --- | --- |
| **CalMS21 Dataset** |  |  |  |  |
| ConstrastivePose | 0.82 | 0.72 | **0.75** | 0.88 |
| No pre-training | 0.79 | 0.72 | 0.72 | 0.88 |

| In-house Dataset |  |  |  |  |
| --- | --- | --- | --- | --- |
| ContrastivePose + overlap in bounding boxes | 0.30 | 0.44 | **0.32** | 0.72 |
| No pre-training + overlap in bounding boxes | 0.28 | 0.40 | 0.29 | 0.68 |

Our method can be used to study the behavior of mice and other animal models, and their social interactions in a setting that allows for free interaction. We demonstrated individual specificity in the data output using our model, which is critical for studies requiring individual-based identification, like research on models of social behavior disorders (e.g. autism spectrum disorders, anxiety disorders). Our method is not limited to any particular type of pose estimation (key points, bounding boxes etc.) or set of behaviors. It can be easily applied for pose estimation and additional behaviors not discussed in this study. This adds to the adaptability of our model to suit various research needs, thereby achieving our goal of existing supervised learning workflow to intelligently automate a task that has high human dependency.

Future work can seek to investigate and incorporate other techniques of self-supervised learning such as pre-text task learning, for e.g. predicting missing values or predicting video clip order, to improve the learnt representation further. The contrastive method proposed in this paper only performs spatial augmentation and thus may not be very effective in extracting useful temporal features. Hence, temporal based tasks may be especially useful for behaviors that happen over a period of time such as one animal following another.

## Materials and Methods

### Datasets

The first dataset is Task 1 of the Caltech Mouse Social Interactions (CalMS21) Dataset[14]. The dataset consists of 7 labeled key points (nose, left ear, right ear, neck, left hip, right hip, and tail-base) for two interacting mice in a box. For each key point, there is the x and y pixel positions. Each frame is labeled for 4 behaviors: attack, investigation, mount and others (non-interaction). Please refer to [14] for details on the dataset.

The second dataset Is video recording from an in-house experiment conducted on two interacting mice in a box. The pose of mice in the video is labeled using the YOLOv3 algorithm[20]. YOLO generated pose-estimation data consists of 4 bounding boxes capturing the nose, head, body, and tail-base for each mouse. For each bounding box, there is the x and y pixel positions of the top left corner of the box, and the height and width of the box. The videos are labeled for 10 behaviors: nose-nose sniff, body sniff 1, body sniff 2, anogenital sniff 1, anogenital sniff 2, mutual circle, affiliative, following 1, following 2 and exploration (behaviors with suffixes 1 and 2 indicate the identity of mice performing the action). Illustrations of behaviors are provided in S2 Table. The use of bounding box tracking by YOLO instead of key points also serves to demonstrate generalizability to different types of pose-estimation methods.

The pose estimation for both dataset are illustrated **Fig *2***.

**Fig 2.**
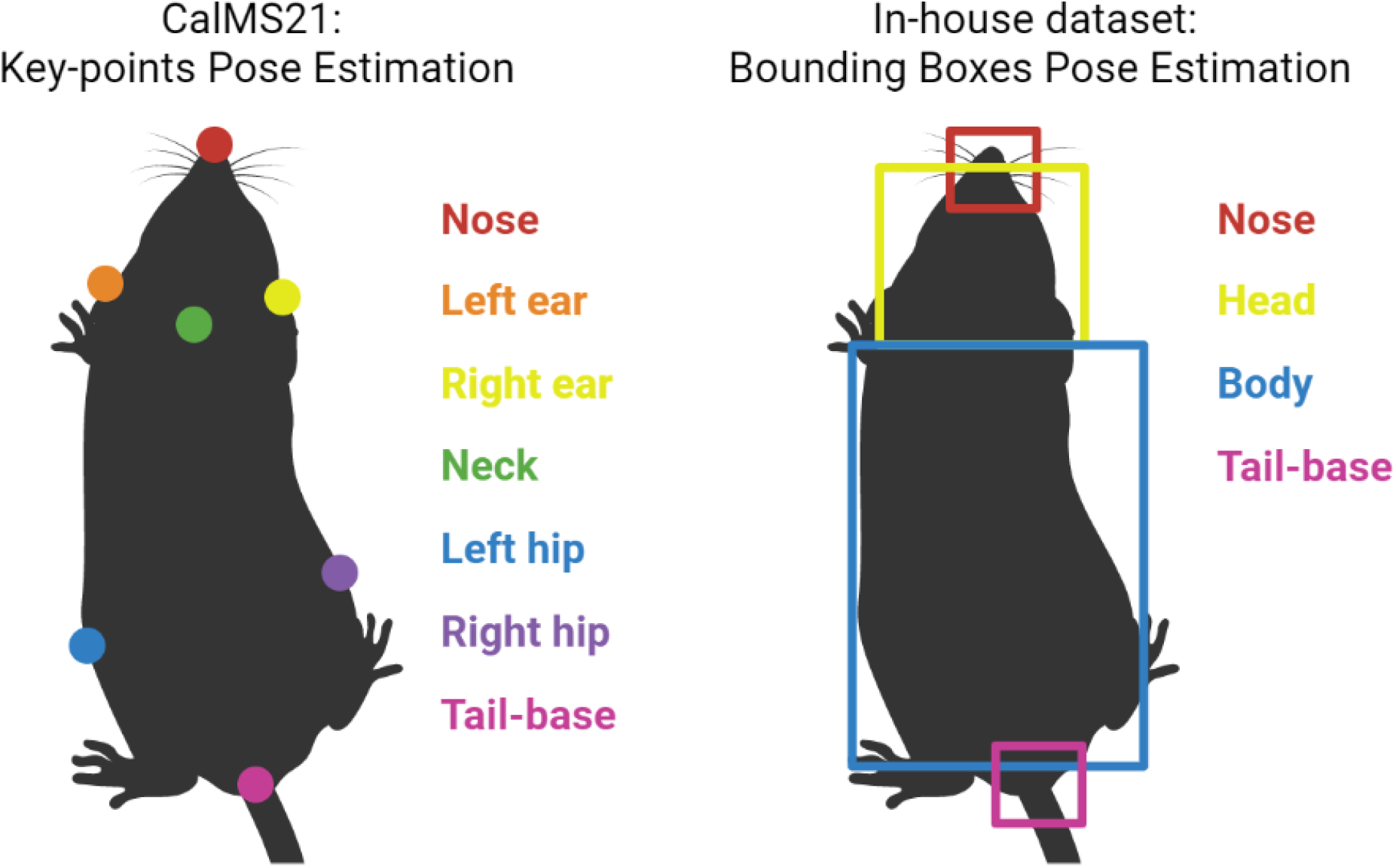
Pose estimation of mice used in CalMS21 and in-house dataset. CalMS21 dataset uses 7 key points, each defined by a x, y positional value. The in-house dataset uses 4 bounding boxes, each defined by the x, y positional value of the top left corner, and the height and width of the box.

### Overview of methods

ConstrastivePose takes as input pose estimation data and outputs a feature representation of the data, which can then be used for downstream supervised classification. It is akin conceptually to the feature engineering step.

The training for ConstrastivePose model uses a large set of unlabeled training data. After training with the unlabeled data through contrastive learning, the model would then be fine-tuned with a relatively small set of labeled training data in a supervised fashion. For the downstream task of supervised classification of behaviors, the small set of labeled training data would be used to train a random forest model (other type of machine learning model may also be used instead). The random forest model takes the learnt representations as computed by the previously trained ConstrastivePose as input. The process is illustrated in **Fig *3***.

**Fig 3.**
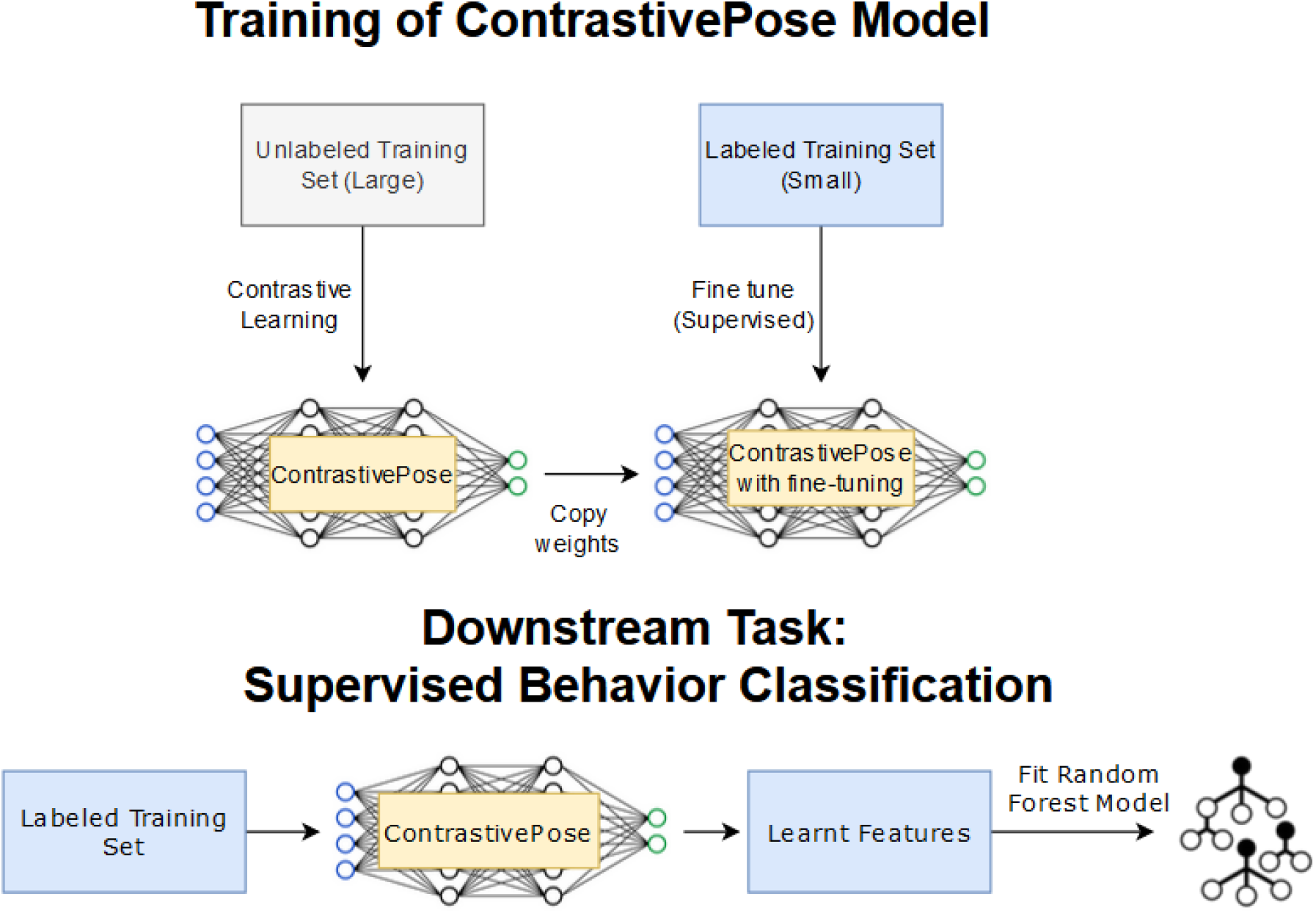
Overview of ConstrastivePose method. The ConstrastivePose model is trained on large set of unlabeled data through contrastive learning, and then fine-tuned on a small set of labeled data. When applying to downstream supervised behavior classification task, the labeled training set is passed to the ConstrastivePose model which outputs the learnt features that can be then used to fit a classifier such as random forest

For inference, we simply take the video that we wish to be labeled, obtain the pose estimation for each frame of the video, pass it through the trained ConstrastivePose model and use its output representation as the input for the trained random forest classifier to obtain the behavior predictions.

The specific details of the training and testing workflow for the data used in this paper are presented in S3.

### Sliding window for input data

To capture temporal aspects of the animal poses which are important for behaviors that take place over many frames, such as following, and attack, we used a sliding window approach to generate the input data for the training and test sets. The length of the sliding window is a hyperparameter. We set this at 30 frames for a 30-frames per second video based on visual inspection that temporal activities can be identified within a second of the video. This hyperparameter has not been tuned. Each datapoint is therefore a matrix of size *30 × number of original features*. For example, for CalMS21, there are 2 mice, 7 key points for each mouse, and 2-D coordinates for each point. We take the frame *t – 29* to frame *t* and concatenate into a *30 × 28* matrix as *X*_*t*_, and the label *Y*_*t*_ will be the behavior labeled for frame *t*.

### Contrastive learning

Contrastive learning has been the most successful self-supervised learning technique used in computer vision, achieving state-of-the-arts performance. Self-supervised learning is an approach to learn from unlabeled samples by generating tasks or pseudo-labels from the data and training a neural network to learn to solve those tasks or pseudo-labels. Some examples of tasks include inpainting missing sections of images or unscrambling scrambled images[23,24].Through this self-supervised training, the model can learn a representation of the data that is also helpful for other downstream tasks. The model can then be fine-tuned with small amounts of training data to be optimized for downstream tasks. Recently, contrastive learning has been applied for feature extraction from animal videos by Jia et al.[25], by performing contrastive learning on the frame images directly in similar fashion to existing work in computer vision.

Contrastive learning’s goal is to learn representations of data such that similar datapoints are close to each other, while dissimilar ones are far apart, without the need for labels[24]. This is achieved during training by using data augmentation. The data augmentation should not change the fundamental characteristic of the data that is relevant for the task at hand. For example, data augmentation in contrastive learning for image classification tasks include random crops and rotations. These augmentations do not change the fundamental characteristic of the data for image classification because a rotated or cropped image still represents the same class of object[24,26].

For pose estimation data, augmentation is achieved by random flipping, rotation and translation of the poses, which do not change the fundamental characteristic for behavior classification. It is the same behavior no matter how we mirror, rotate, or translate the setup. Hence, we can define a data augmentation that performs random flipping along the *x* or *y* axis, rotation by random angles, and random translation along both axes. For details on the implementation of augmentation, please refer to Supplementary Materials.

During training, the model learns to reduce the difference in representations between any image and its random augmented version and enlarge the difference with other images in the batch[24,26]. The training process is illustrated in **Fig *4***. For detailed steps of the implementation of contrastive learning, and neural network architecture used, refer to S4 – S6.

**Fig 4.**
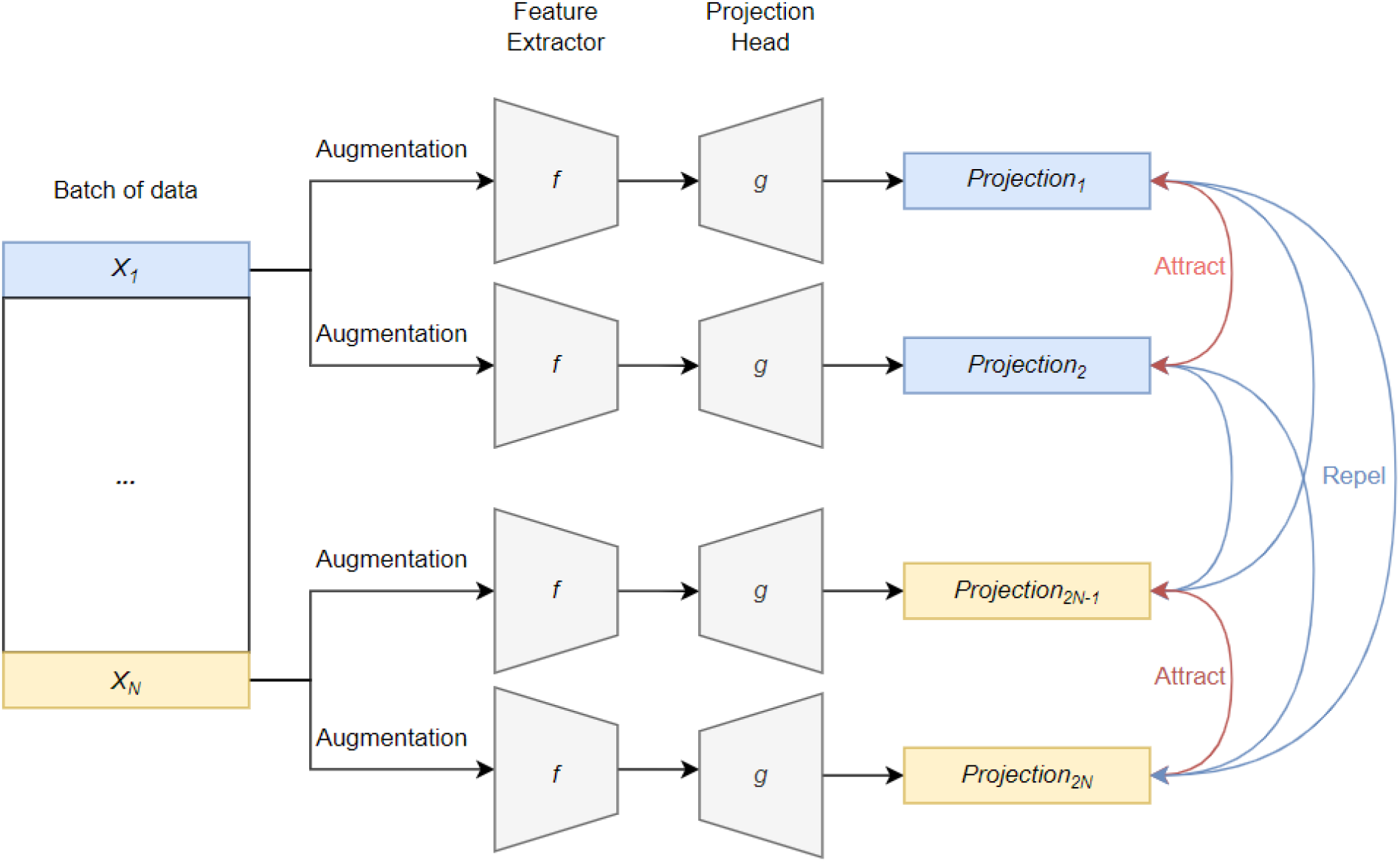
Contrastive learning process. For a given batch of data, each data point goes through parallel random augmentation and gets passed through the networks to obtain two projections. The contrastive loss is computed over the whole batch. By minimizing the contrastive loss, the model seeks to make the matching pairs’ projections more similar while making all other pairs’ projections dissimilar

### Feature engineering methods

We describe the methods used to compute the engineered features used as comparison in this paper.

Velocity features are computed by the difference position between any frame and its previous frame. For example, in the CalMS21 dataset, with 2 mice each with 7 body parts described by *x, y* coordinates, there are 28 velocity features.

Distances between key points are computed as the Euclidean distance between the combination of any key point of one mouse with any key point of the other mouse. For example, in the CalMS21 dataset, there are 7 × 7 key point pairs, which give 49 pairwise distance features.

For bounding boxes, the overlap in bounding boxes is computed as the intersection area between two bounding boxes, divided by total area covered by both boxes. Similar to pairwise distances, the overlap ratio is computed for all combinations of key points of one mouse with key points of the other mouse. For example, in our in-house dataset, there are 4 × 4 key point pairs giving 16 pairwise overlap ratio features.

## Acknowledgments

N/A

## References

1. von Ziegler L, Sturman O, Bohacek J. Big behavior: challenges and opportunities in a new era of deep behavior profiling. Neuropsychopharmacology. 2021;46: 33–44. doi:10.1038/s41386-020-0751-7

2. Bohlen M, Hayes ER, Bohlen B, Bailoo JD, Crabbe JC, Wahlsten D. Experimenter effects on behavioral test scores of eight inbred mouse strains under the influence of ethanol. Behav Brain Res. 2014;272: 46–54. doi:10.1016/j.bbr.2014.06.017

3. Garcia VA, Junior CFC, Marino-Neto J. Assessment of observers’ stability and reliability — A tool for evaluation of intra- and inter-concordance in animal behavioral recordings. Annu Int Conf IEEE Eng Med Biol Soc 2010. 2010. pp. 6603–6606. doi:10.1109/IEMBS.2010.5627131

4. Mathis A, Mamidanna P, Cury KM, Abe T, Murthy VN, Mathis MW, et al. DeepLabCut: markerless pose estimation of user-defined body parts with deep learning. Nat Neurosci. 2018;21: 1281–1289. doi:10.1038/s41593-018-0209-y

5. Sturman O, von Ziegler L, Schläppi C, Akyol F, Privitera M, Slominski D, et al. Deep learning-based behavioral analysis reaches human accuracy and is capable of outperforming commercial solutions. Neuropsychopharmacology. 2020;45: 1942–1952. doi:10.1038/s41386-020-0776-y

6. van den Boom BJG, Pavlidi P, Wolf CJH, Mooij AH, Willuhn I. Automated classification of self-grooming in mice using open-source software. J Neurosci Methods. 2017;289: 48–56. doi:10.1016/j.jneumeth.2017.05.026

7. Bailoo JD, Bohlen MO, Wahlsten D. The precision of video and photocell tracking systems and the elimination of tracking errors with infrared backlighting. J Neurosci Methods. 2010;188: 45–52. doi:10.1016/j.jneumeth.2010.01.035

8. Geuther BQ, Deats SP, Fox KJ, Murray SA, Braun RE, White JK, et al. Robust mouse tracking in complex environments using neural networks. Commun Biol. 2019;2: 1–11. doi:10.1038/s42003-019-0362-1

9. Sturman O, Germain PL, Bohacek J. Exploratory rearing: a context- and stress-sensitive behavior recorded in the open-field test. Stress. 2018;21: 443–452. doi:10.1080/10253890.2018.1438405

10. Wu S, Tan KJ, Govindarajan LN, Stewart JC, Gu L, Ho JWH, et al. Fully automated leg tracking of drosophila neurodegeneration models reveals distinct conserved movement signatures. PLoS Biol. 2019;17: e3000346. doi:10.1371/journal.pbio.3000346

11. Hong W, Kennedy A, Burgos-Artizzu XP, Zelikowsky M, Navonne SG, Perona P, et al. Automated measurement of mouse social behaviors using depth sensing, video tracking, and machine learning. Proc Natl Acad Sci U S A. 2015;112: E5351–E5360. doi:10.1073/pnas.1515982112

12. Nilsson SR, Goodwin NL, Choong JJ, Hwang S, Wright HR, Norville ZC, et al. Simple Behavioral Analysis (SimBA) – an open source toolkit for computer classification of complex social behaviors in experimental animals. BioRxiv [Preprint]. 2020 [cited 22 Oct 2022]. doi:10.1101/2020.04.19.049452

13. Lorbach M, Kyriakou EI, Poppe R, van Dam EA, Noldus LPJJ, Veltkamp RC. Learning to recognize rat social behavior: Novel dataset and cross-dataset application. J Neurosci Methods. 2018;300: 166–172. doi:10.1016/j.jneumeth.2017.05.006

14. Sun JJ, Karigo T, Chakraborty D, Mohanty SP, Wild B, Sun Q, et al. The Multi-Agent Behavior Dataset: Mouse Dyadic Social Interactions. arXiv [Preprint]. 2021 [cited 22 Oct 2022]. doi:10.48550/arXiv.2104.02710

15. Segalin C, Williams J, Karigo T, Hui M, Zelikowsky M, Sun JJ, et al. The mouse action recognition system (MARS) software pipeline for automated analysis of social behaviors in mice. Elife. 2021;10. doi:10.7554/eLife.63720

16. Hsu AI, Yttri EA. B-SOiD, an open-source unsupervised algorithm for identification and fast prediction of behaviors. Nat Commun. 2021;12. doi:10.1038/s41467-021-25420-x

17. Batpurev T, Shibata T, Matsumoto J, Nishijo H. Automatic Identification of Mice Social Behavior Through Multi-Modal Latent Space Clustering. 2021 Joint 10th International Conference on Informatics, Electronics and Vision, ICIEV 2021 and 2021 5th International Conference on Imaging, Vision and Pattern Recognition, icIVPR 2021. Institute of Electrical and Electronics Engineers Inc.; 2021. doi:10.1109/ICIEVICIVPR52578.2021.9564213

18. Arac A, Zhao P, Dobkin BH, Carmichael ST, Golshani P. DeepBehavior: A Deep Learning Toolbox for Automated Analysis of Animal and Human Behavior Imaging Data. Front Syst Neurosci. 2019;13: 20. doi:10.3389/fnsys.2019.00020

19. Graving JM, Chae D, Naik H, Li L, Koger B, Costelloe BR, et al. Deepposekit, a software toolkit for fast and robust animal pose estimation using deep learning. Elife. 2019;8. doi:10.7554/eLife.47994

20. Redmon J, Divvala S, Girshick R, Farhadi A. You only look once: Unified, real-time object detection. Proc IEEE Comput Soc Conf Comput Vis Pattern Recognit. 2016;2016-December: 779–788. doi:10.1109/CVPR.2016.91

21. Jhuang H, Garrote E, Yu X, Khilnani V, Poggio T, Steele AD, et al. Automated home-cage behavioural phenotyping of mice. Nat Commun. 2010;1: 1–10. doi:10.1038/ncomms1064

22. Rousseau JBI, van Lochem PBA, Gispen WH, Spruijt BM. Classification of rat behavior with an image-processing method and a neural network. Behav Res Methods Instrum Comput. 2000;32: 63–71. doi:10.3758/BF03200789

23. Doersch C, Zisserman A. Multi-task Self-Supervised Visual Learning. Proc IEEE Int Conf Comput Vis. Institute of Electrical and Electronics Engineers Inc.; 2017. pp. 2070–2079. doi:10.1109/ICCV.2017.226

24. Khan A, Albarri S, Manzoor MA. Contrastive Self-Supervised Learning: A Survey on Different Architectures. 2nd IEEE International Conference on Artificial Intelligence, ICAI 2022. 2022; 1–6. doi:10.1109/ICAI55435.2022.9773725

25. Jia Y, Li S, Guo X, Lei B, Hu J, Xu XH, et al. Selfee, Self-supervised Features Extraction of animal behaviors. Elife. 2022;11. doi:10.7554/ELIFE.76218

26. Chen T, Kornblith S, Norouzi M, Hinton G. A Simple Framework for Contrastive Learning of Visual Representations. arXiv [Preprint]. 2020 [cited 22 Oct 2022]. doi:10.48550/arXiv.2002.05709

27. Luxem K, Mocellin P, Fuhrmann F, Kürsch J, Remy S, Bauer P. Identifying Behavioral Structure from Deep Variational Embeddings of Animal Motion. BioRxiv [Preprint]. 2022. doi:10.1101/2020.05.14.095430

